# The effects of training population design on genomic prediction accuracy in wheat

**DOI:** 10.1101/443267

**Authors:** Stefan McKinnon Edwards, Jaap B. Buntjer, Robert Jackson, Alison R. Bentley, Jacob Lage, Ed Byrne, Chris Burt, Peter Jack, Simon Berry, Edward Flatman, Bruno Poupard, Stephen Smith, Charlotte Hayes, R. Chris Gaynor, Gregor Gorjanc, Phil Howell, Eric Ober, Ian J. Mackay, John M. Hickey

## Abstract

Genomic selection offers several routes for increasing genetic gain or efficiency of plant breeding programs. In various species of livestock there is empirical evidence of increased rates of genetic gain from the use of genomic selection to target different aspects of the breeder’s equation. Accurate predictions of genomic breeding value are central to this and the design of training sets is in turn central to achieving sufficient levels of accuracy. In summary, small numbers of close relatives and very large numbers of distant relatives are expected to enable accurate predictions.

To quantify the effect of some of the properties of training sets on the accuracy of genomic selection in crops we performed an extensive field-based winter wheat trial. In summary, this trial involved the construction of 44 F_2:4_ bi- and triparental populations, from which 2992 lines were grown on four field locations and yield was measured. For each line, genotype data were generated for 25,000 segregating single nucleotide polymorphism markers. The overall heritability of yield was estimated to 0.65, and estimates within individual families ranged between 0.10 and 0.85. Within cross genomic prediction accuracies of yield BLUEs were 0.125 – 0.127 using two different cross-validation approaches, and generally increased with training set size. Using related crosses in training and validation sets generally resulted in higher prediction accuracies than using unrelated crosses. The results of this study emphasize the importance of the training set design in relation to the genetic material to which the resulting prediction model is to be applied.

## Introduction

Genomic selection in plant breeding offers several routes for increasing the genetic gain or efficiency of plant breeding programs (e.g., Bernardo and Yu, 2007; Hickey et al., 2014; Gaynor et al., 2017). Genomic selection based strategies can achieve this by reducing breeding cycle time, increasing selection accuracy and increasing selection intensity; three of the four factors in the breeder’s equation. Genomic prediction can reduce breeding cycle time because individuals can be selected and crossed without being phenotyped. It can increase the selection accuracy because genomic data enables more powerful statistical models and experimental designs using more observations than can be phenotyped in a single trial round. By reducing the cost of evaluating individuals via reducing the numbers phenotyped and/or reducing their replication, application of genomic selection can increase selection intensity. A final advantage is that the prediction models may be cumulatively updated with data of trials from previous years and become more accurate, enabling individuals to be “evaluated” across a broader range of environments and years.

In livestock there is empirical evidence of increased rates of genetic gain from the use of genomic selection to target different aspects of the breeder’s equation. For example the first seven years of genomic selection in US dairy cattle has delivered ~50 - 100% increases in rates of genetic gain (García-Ruiz et al., 2016). Much of this gain has emanated from a reduction in generation interval. In commercial pig breeding, genomic selection has driven a 35% increase in rate of genetic gain in the breeding program that supplies the genetics in 25% of the intensively raised pigs globally. This gain came from increased accuracy of selection and a better alignment of selection accuracy with the breeding goal (W. Herring, personal communication).

Genomic selection uses genotype data to calculate the realised relationship between individuals, and in a standardized statistical framework uses data from phenotyped relatives to estimate genetic values of the selection candidates. The usefulness of genomic selection to a breeder is a function of its accuracy. This is affected by the relatedness between the phenotyped individuals in the training set and the individuals that are to be predicted (Habier et al., 2007, 2010; Meuwissen, 2009; Clark et al., 2012; Hickey et al., 2014; Liu et al., 2016), which may or may not be phenotyped themselves. In addition to the level of relatedness, the sample size of the phenotyped individuals is an important factor in determining accuracy (Zhang et al., 2017).

In summary, small numbers of close relatives and very large numbers of distant relatives enable accurate predictions. Small or modest numbers of distant relatives do not enable accurate predictions, as they share only a small proportion of genome with the selection candidates, and thus provide less reliable predictions (de los Campos et al., 2013). Finally, the training set should also comprise a diverse set of individuals to produce reliable predictions (Calus, 2010; Pszczola et al., 2012; Pszczola and Calus, 2015), as supported by recent research in both cattle (Jenko et al., 2017) and simulated barley (Neyhart et al., 2017).

The objective of this study was to explore the effect of level of relatedness between training set and validation set on genomic prediction accuracy using data from a large set of field experiments. To do this, 44 bi-parental or three-way crosses were obtained from four commercial wheat breeders in the United Kingdom, as described for the GplusE Project (Mackay et al., 2015). The crosses had different degrees of relatedness among each other and there were many shared parents. 68 F_2:4_ lines from each cross were genotyped and phenotyped for yield. As this data set is of substantial size, it enabled genomic predictions while masking specific fractions to assess the impact on genomic selection accuracy of training sets: (i) of different sizes; and (ii) that comprise close or distant relatives, or combinations thereof.

## Materials and Methods

### Germplasm

Thirty-nine bi-parental and 5 triparental populations were used to develop 2992 F_2:4_ lines (68 per cross). The parents of these populations were elite breeders’ germplasm consisting of both hard and soft winter wheat cultivars adapted to the United Kingdom. A total of 27 parents were used, of which 5 parents were used in 6 or more crosses, 6 parents were used in 3 or 4 crosses, and 1 parent was used in 2 crosses. The remaining 15 parents were only used in a single cross.

### Genotypes

The F_2:4_ lines were genotyped using the Wheat Breeders’ 35K Axiom array (Allen et al., 2016). The DNA for genotyping was obtained by bulking leaves from approximately 6 F_4_ plants per F_2:4_ line. Genotype calling was performed using the Axiom Analysis Suite 2.0 with a modified version of the “best practices” workflow. Quality control threshold was reduced to 95 (97 normally), plate pass percent was changed to 90 (95 normally), and average call rate was changed to 97 (98.5 normally). After quality control and genotype calling, a total of 35,143 markers were brought forward with 24,498 segregating in the 44 crosses.

### Phenotypes

The F_2:4_ lines and agronomic checks were evaluated in 2 by 4 meter harvested plots at 2 locations (Cambridge, UK and Duxford, UK) in the 2015-16 growing season, and 2 locations (Hinxton, UK, and Duxford, UK) in the 2016-17 growing season. All locations were managed for optimal yield by following best agronomic practice. All F_2:4_ lines were evaluated in 4 plots. Seed for eleven of the populations was unavailable in the 2015-16 growing season. To accommodate these populations and keep the number of plots per line constant, an allocation of F_2:4_ lines was devised that was highly unbalanced across both years and locations as described below.

In the 2015-16 growing season, 33 of the 44 populations were planted at two locations (Table 1). The experimental design for both locations was a modified α-lattice design (Patterson and Williams, 1976). The design consisted of a traditional, replicated α-lattice design with un-replicated lines added to the sub-blocks. The replicated portion of the alpha-lattice design was composed of the agronomic checks and half of the lines (34) from 22 of the F_2:4_ populations. These lines were planted in 2 blocks split into 151 sub-blocks each containing 5 lines. The remaining F_2:4_ lines were randomly allocated to sub-blocks, bringing the total number of lines per sub-block to either 9 or 10. Half of the F_2:4_ lines used for the replicated portion of the design differed between locations. Thus lines from 22 of the F_2:4_ populations were evaluated in 3 plots split across both locations and the lines from the remaining populations were evaluated in 2 plots split across locations.

**Table 1:** Trial design summary showing number of plots per tested line per location.

| # Lines | 2015/2016 |  | 2016/2017 |  |
| --- | --- | --- | --- | --- |
|  | Cambridge | Duxford | Duxford | Hinxton |
| 367 | 2 | 1 | 1 | 0 |
| 381 | 2 | 1 | 0 | 1 |
| 381 | 1 | 2 | 1 | 0 |
| 367 | 1 | 2 | 0 | 1 |
| 748 | 1 | 1 | 1 | 1 |
| 748 | 0 | 0 | 2 | 2 |
| **Total plots** | **2992** | **2992** | **2992** | **2992** |

All 44 populations were planted in the 2016-17 growing season at two locations (Table 1); the experimental design was similar as in the previous season. The replicated portion of the α-lattice design was composed of the agronomic checks and the F_2:4_ lines from the 11 populations not planted in the 2015-16 growing season. These lines were planted in 2 blocks split into 156 sub-blocks each containing 5 lines. Additional F_2:4_ lines from the other populations were randomly allocated to sub-blocks, bring the total number of lines per sub-block to 10.

### Yield Trial Analysis

Yield phenotypes were spatially adjusted for each trial separately. An AR1 x AR1 model (Gilmour et al., 1997) was used to adjust spatial variation across both columns and rows as implemented in ASREML 3.0.22 (Gilmour et al., 2009). A summary of line means after adjusting for spatial effects is shown in Table 2.

**Table 2:** Summary of line means per location after adjusting for spatial effects.

|  |  | No. lines | Avg. value | Coef. Variation | Correlation<sup>†</sup> |
| --- | --- | --- | --- | --- | --- |
| 2016 | Cambridge | 2,247 | 8.58 | 6.1% | 0.63 |
| 2016 | Duxford | 2,248 | 10.82 | 6.3% | 0.81 |
| 2017 | Hinxton | 2,249 | 4.64 | 10.3% | 0.71 |
| 2017 | Duxford | 2,235 | 8.24 | 6.6% | 0.62 |

<sup>†</sup>: Correlation between moisture corrected yield values and spatially adjusted values.

Best linear unbiased estimates (BLUEs) for each line were estimated collectively across all trials by fitting the following model:

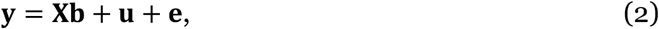

where **y** was the response vector of spatially adjusted yield values, **b** site-specific means with design matrix **X**, **u** line BLUEs to estimate, and **e** the model residual.

### Genomic prediction

This study used the genomic best linear unbiased prediction (GBLUP) model to estimate heritabilities and predict line effects. The GBLUP model used was:

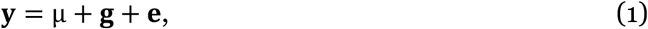

where **y** was the response vector of yield BLUEs, μ the model intercept, **g** the vector of genetic values of genotyped F_2:4_ and **e** the model residual. We assumed that 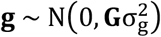 with genomic relationship matrix calculated as **G** = **WW**′/2∑p_i_(1 – p_i_) (VanRaden, 2008) from the centred genotype matrix **W** and allele frequencies p_i_ estimated in the dataset. Further, we assumed that 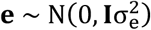, which was assumed uncorrelated to **g**.

The Average-Information Restricted Maximum Likelihood (AI-REML) algorithm (Madsen et al., 1994; Johnson and Thompson, 1995), as implemented in DMU v. 5.1 (Madsen and Jensen, 2000), was used to fit the GBLUP model to a subset of the data (training set) and predict line effects (ĝ) in the validation set. We defined convergence of the AI-REML algorithm based on the change of variance components, |*θ*^(*t*+1)^ − *θ*^(*t*)^| < 10^−5^, where *θ*^(*t*)^ is the vector of normalised variance components estimated at step K (Jensen et al., 1997).

The heritability was calculated from the trial yield data per plot as 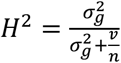 in which n is the number of locations in which the genotype was observed and v is the residual variance (Piepho and Mohring, 2007).

### Prediction accuracies

We applied several cross-validation strategies for investigating prediction accuracies of genomic selection with varying training set size and grouping of training sets and validation sets, as described in detail in the following sections. In all strategies, the GBLUP model was used as described above. The prediction accuracies were calculated as the Pearson correlation (ρ) between the yield BLUEs and its prediction from the GBLUP model.

#### Cross-validation prediction accuracy

In the first approach, we used 10-fold cross-validation and leave-one-cross-out cross-validation (effectively 44-fold cross-validation; refer to Figure 1). Populations were randomly assigned to either training or validation set, without considering that some crosses are more closely related due to sharing a parent or other ancestors. The validation sets were entire populations, which means that line means of a population was confined entirely to either training set or validation set. Prediction accuracies were summarised on a per cross basis and encapsulate the within cross genomic prediction accuracy (sometimes referred to as the within family accuracy or the accuracy of predicting the Mendelian sampling term). For the 10-fold cross-validation, 10 replicates were performed where the 10 folds were re-sampled.

**Figure 1:**
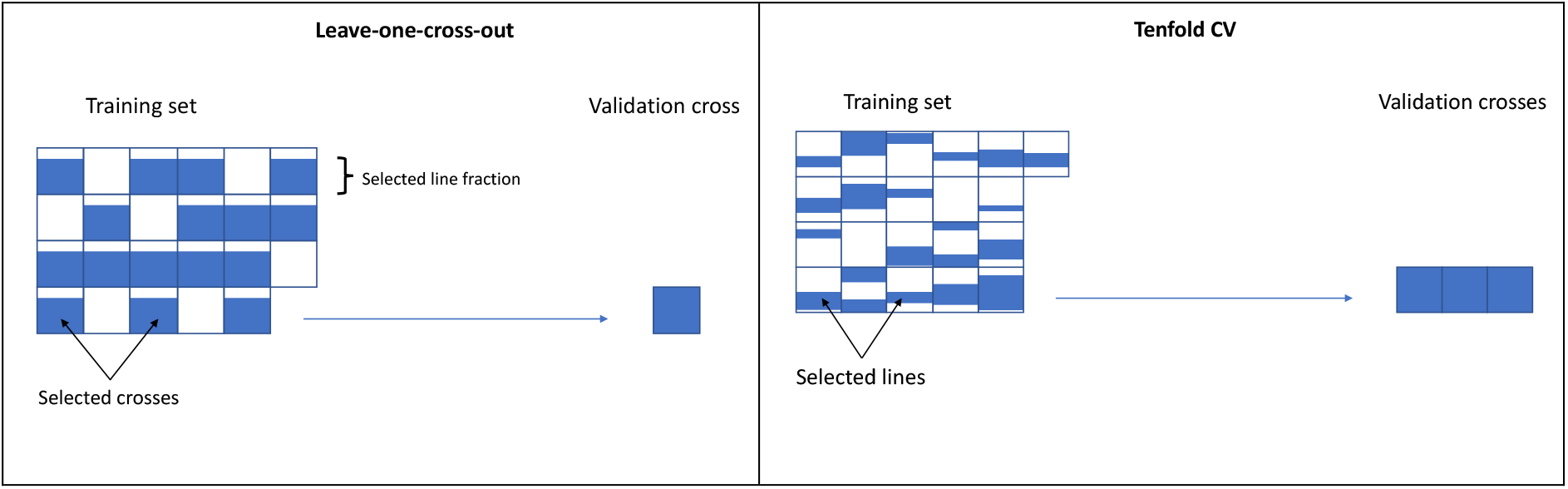
Resampling strategies applied to assess the impact of training set design. Leave-one-cross out strategy (left) tests the impact of inclusion of the amount of crosses as well as training set size, while the ten-fold cross validation (right) tests training set size only.

To evaluate the effect of training set size, the two cross-validation methods described above were repeated using a subset of the total training set. For the 10-fold cross-validation, 10%, 20%, …, 80%, 90% of records were randomly removed from the training set, before estimating variance components and predicting line means of the validation set. For each replicate and the proportion of training set masked, 10 repetitions were performed and the resulting prediction accuracies encapsulate the joint across and within cross genomic prediction accuracy. For the leave-one-cross-out cross-validation, 1 – 10, 15, 20, 30, 40 crosses were randomly sampled to be used as training set. For each number of crosses sampled as training sets, 10, 20, …, 60, 65 records from each cross was sampled. Again, 10 repetitions were performed. We emphasise that the validation sets were always entire populations (from 3-4 crosses in 10-fold cross-validations, from single cross in leave-one-cross-out) and no records of the validated populations were included in the training set.

#### Prediction accuracy with related or unrelated crosses

In the second approach, we evaluated the prediction accuracies under different levels of relatedness between validation and training sets. The 6 crosses of the 4 most frequently used parents were targeted as validation crosses and tested separately. In summary, the training sets consisted of varying proportions of sister-lines and half-sibs from offspring of either one or both parents or unrelated crosses. Specifically, for each validation cross, training sets were designed to consist of either one or several crosses of one parent, an equal number of crosses from each parent, nominally unrelated crosses, or equal number of related and unrelated crosses. To reduce computation time, for each training set of crosses, 5 combinations were sampled from the large number of possible combinations. For each training set, the validation cross contributed with 0, 1, 2, or 3 quarters of its lines. The prediction accuracies were evaluated for the fourth quarter of lines that were not used in the training set. For each combination of training set, 10 replicates were performed as well as cycling through all four quarters of the validation cross as training set.

### Results

44 bi- and tri-parental crosses from 27 parents were analysed for yield with a GBLUP model (1), using BLUEs from 4 trials (2 trials in 2016, and 2 trials in 2017).

#### Trait heritability

The overall heritability of yield for all populations over all four trial locations was estimated at 0.65. Heritabilities estimated on single crosses were highly variable, ranging from as low as 0.1 to as high as 0.85 (Figure 2).

**Figure 2:**
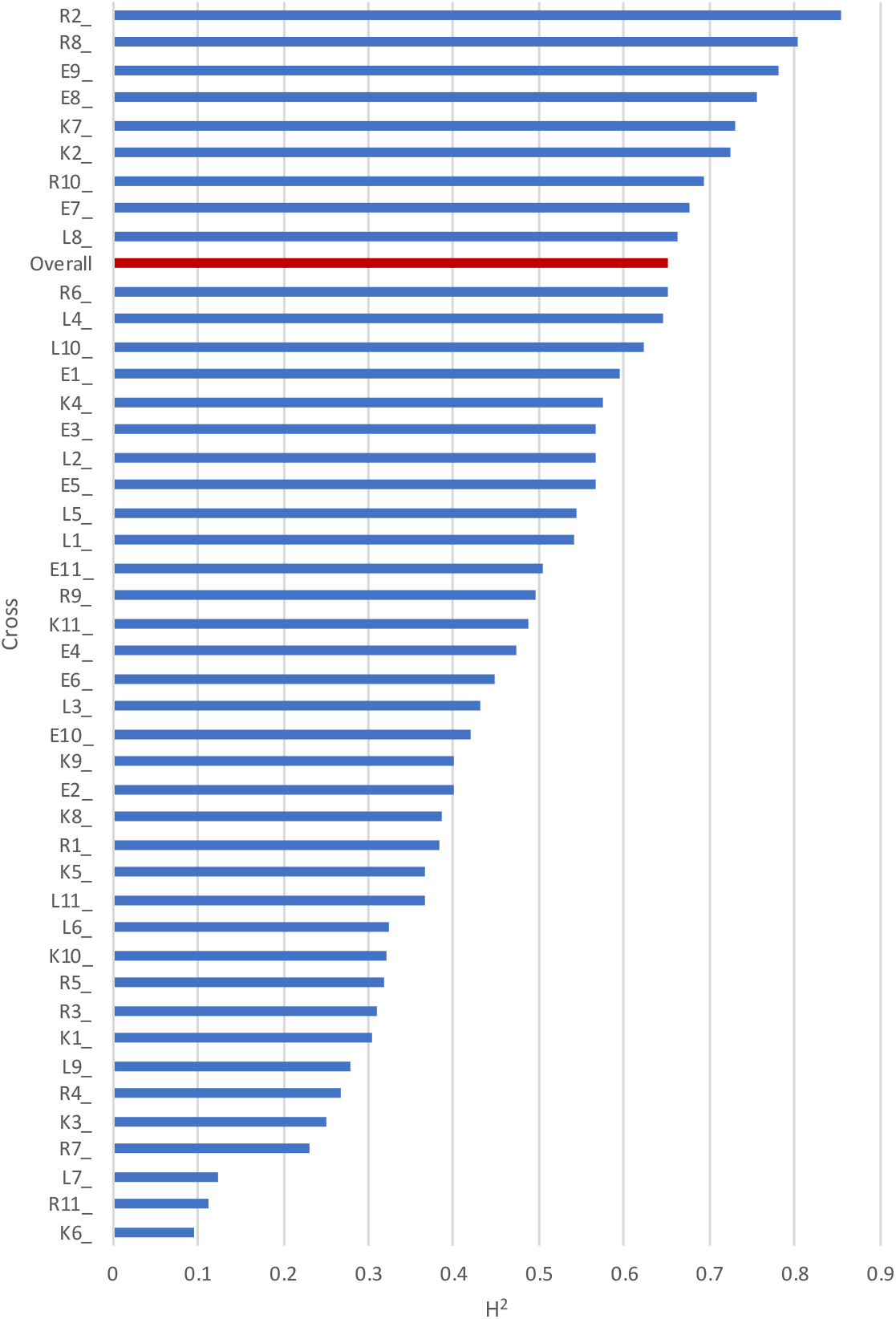
Yield heritabilities when estimated per cross. Crosses (blue bars) are ordered by heritability value, overall heritability for this trait is shown in red.

#### Cross-validation prediction accuracy

Within cross prediction accuracies were 0.125 – 0.127 using two different cross-validation approaches (Table 3). In these two approaches, all lines of the crosses used for validation were absent from the training set. Using a 10-fold cross-validation approach where individual lines, not all lines of a cross, were selected for validation sets, the prediction accuracy was slightly higher (0.142) when calculated on a per-cross basis (’10-fold, random’, Table). Lastly, the prediction accuracy was higher when calculated across all crosses in the validation set, due to capturing variation within and between crosses (0.289 and 0.543, Table 3).

**Table 3:**
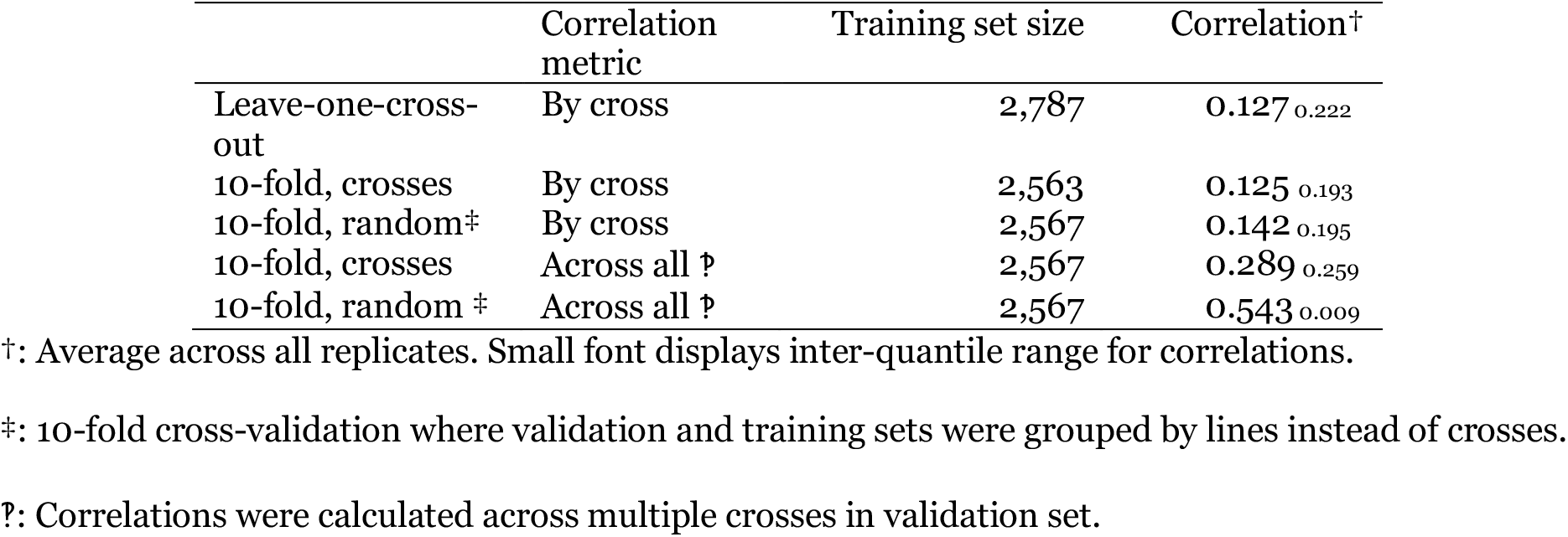
Prediction accuracies using the largest training sets by cross-validation approach.

The prediction accuracy was found to increase with training set size. Figure 3 displays the average prediction accuracy across all crosses with 10^th^ and 90^th^ percentile range shown as the greyed area. The prediction accuracy varied greatly between the crosses (Supplemental figure 1) with some accuracies as high as 0.45 (cross 7), as low as −0.20 (cross 30). For 31 crosses out of 44, significant positive prediction accuracies were found (Wald’s test, p<0.05). Crosses with higher phenotypic variance generally yielded higher predictions; in Supplemental figure 1, prediction accuracy plots for individual crosses are sorted with decreasing phenotypic variance. Finally, the two cross-validation approaches generally produced similar results (Supplemental figure 1), but when the training sets were small, the accuracy of predictions from leave-one-cross-out were less stable than from 10-fold cross validation. The leave-one-cross-out sampled entire crosses in contrast to the 10-fold cross-validation, where lines across all crosses except the validated cross were sampled.

**Figure 3:**
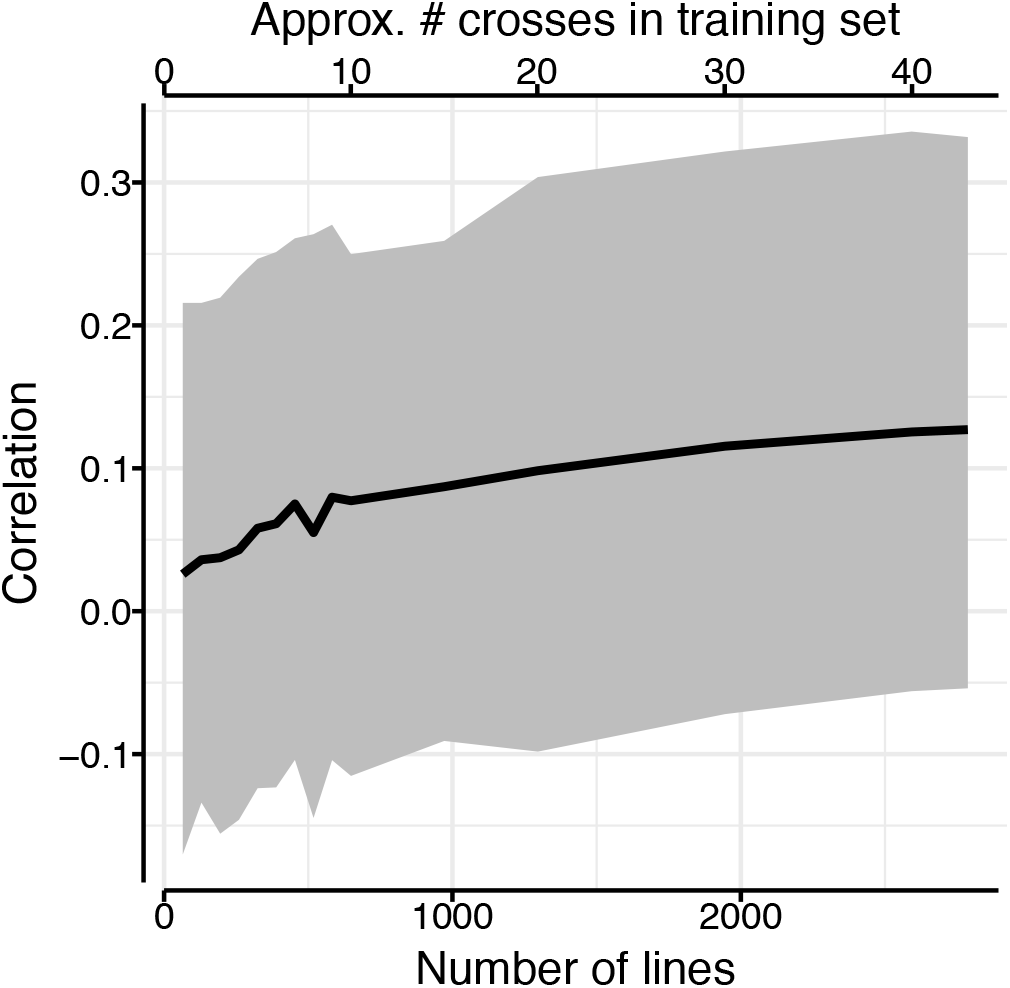
Increasing training set size increased prediction accuracy (correlation). Solid line shows average of all leave-one-cross-out cross-validations with 10^th^ and 90^th^ percentile range shown by greyed area.

The prediction accuracy increased with an increasing number of crosses in training set or increasing number of lines per cross in training set. Figure displays the average prediction accuracy when sampling a number of lines from a number of crosses (x-axis). Adding an additional 10 or 15 lines to a training set of 50 lines per cross generally led to a low increase in prediction accuracy as compared to adding them to training sets of ≤ 40 lines per cross, irrespective of the number of crosses included in the training set.

#### Prediction accuracies with related or unrelated crosses

Using related crosses as a training set generally resulted in higher prediction accuracies compared to using unrelated crosses. This is shown in Figure 5, where the green lines (related training sets) are above the purple lines (unrelated training sets). Using both related and unrelated crosses in equal proportions (blue lines, Figure 5) led generally to similar correlations to those for related crosses. At approximately 7oo to 800 lines in the training set, the prediction accuracy using both related and unrelated crosses plateaued; this was where additional crosses in the training set were unrelated to the validation cross. The level of prediction accuracy of the training set comprising both related and unrelated crosses (lower blue line, Figure) was higher than that in Figure because results in Figure are averages over just 6 crosses rather than over all crosses as in Figure 3.

**Figure 4:**
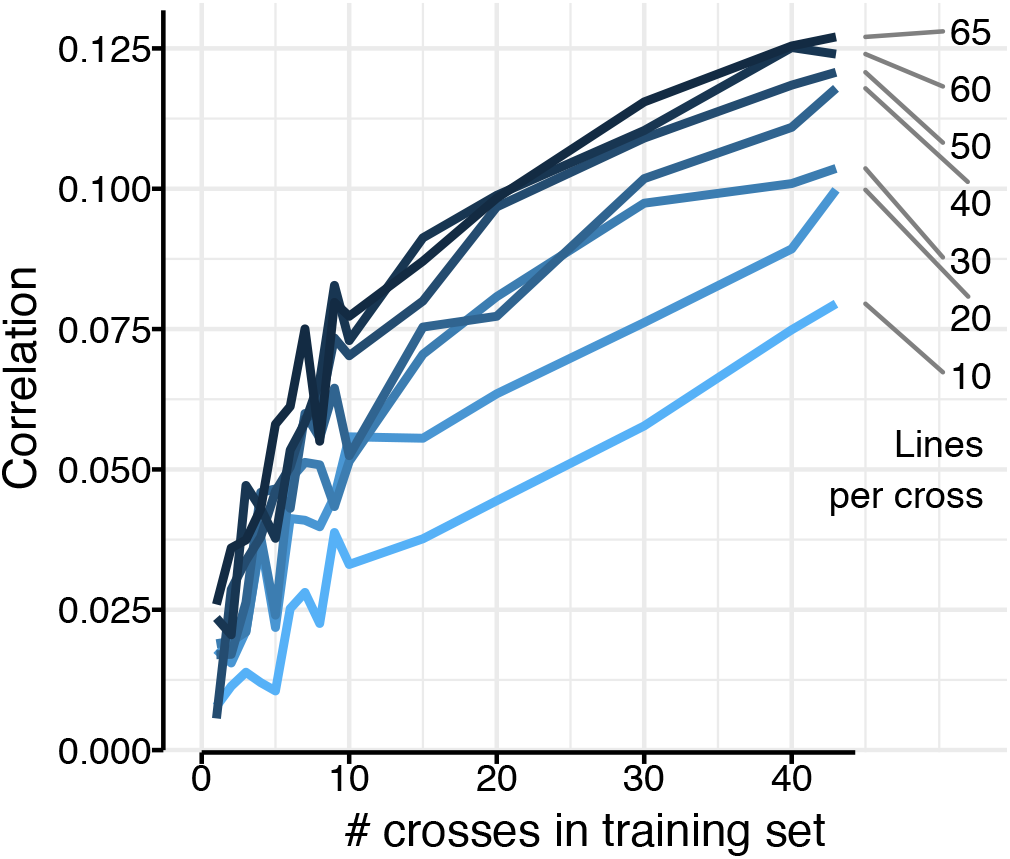
Prediction accuracies increased with the increasing number of crosses or the increasing number of lines per cross in training set. Right-hand numbers show number of lines per cross in training set.

**Figure 5:**
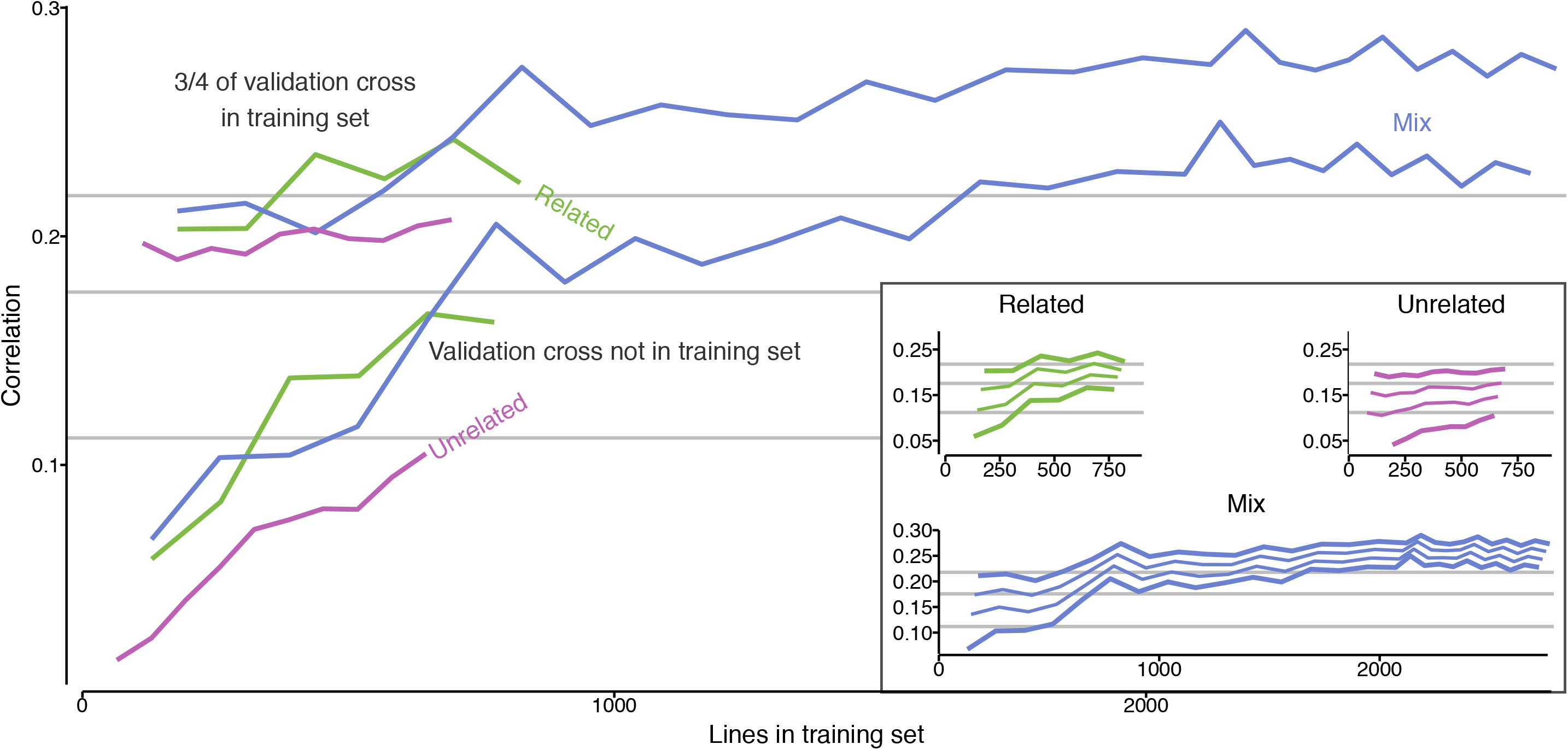
Prediction accuracies increased when the validation cross was partly in training set or had its related crosses in training set. Results show average prediction accuracies for 6 validation crosses. Lines show prediction accuracies when training set comprised of related crosses (green solid line), unrelated crosses (purple dashed line), or a mix of both (blue dotted line). Lower set of lines show prediction accuracies when validation crosses were not included on the training set; upper set of lines show prediction accuracies when validation crosses were included in the training set with 3/4 of lines. Grey horizontal lines show average prediction accuracy using *only* 1/4, 2/4, or 3/4 of validation cross as training set. Inserted figure shows the increase in accuracy when adding 1/4, 2/4, and 3/4 of the validation group to the training set. The thick lines in the inserted figure denote the lines of the main figure.

Using only 1, 2, or 3 quarters of the validation cross as training set (grey, horizontal lines, Figure 5) generally led to prediction accuracies that were higher than using a few unrelated or related crosses as the training set. Adding three quarters of the validation cross to the training sets of other crosses generally increased the prediction accuracy, as shown with the upper thick lines in Figure . The gradual increase in prediction accuracy when adding 1, 2, or 3 quarters of the validation cross to the training set is shown in the inserted plot in Figure 5.

### Discussion

In this study, we have demonstrated the impact of training set size and relatedness on genomic prediction in wheat, using F_2:4_ lines from 44 bi- and tri-parental crosses. The results were consistent with expectations from existing literature (as discussed in the next sections). Specifically, we found that increasing the size of the genomic prediction training set increased accuracy. We also found that training sets composed of lines more closely related to the validation set produce higher prediction accuracies than equivalently sized training sets of more distantly related lines.

It is important for genomic prediction of a complex trait that it displays a reasonable heritability. Our estimate of broad sense heritability for yield (0.65) is well within range of similar studies in wheat (Poland et al., 2012; Combs and Bernardo, 2013; Michel et al., 2016; Schopp et al., 2017; Norman et al., 2017). We note that the heritability values within individual families (Figure) cover the whole range of heritability for this trait reported in the literature.

The various strategies of data subset masking applied in this study has enabled us to demonstrate both training set size and relatedness as parameters that influence successful genomic prediction. Generally, increasing the training set size increased the prediction accuracy, as expected from existing theory (Daetwyler et al, 2008, Goddard, 2009, Hickey et al., 2014) and field reports (Liu et al., 2016; Zhang et al., 2017). However, we can add three observations that put some nuance to this general conclusion. First (1), with a fixed training set size, it is better to increase the number of populations (crosses) rather than number of lines per population (cross). Second (2), the prediction accuracy plateaus when adding additional crosses that are unrelated to the predicted cross (Figure). Third (3), prediction accuracies vary greatly between individual crosses and this could not be explained by neither the crosses’ phenotypic variance nor heritability.

For item 1), we showed that, for example, using 10 crosses with 40 lines per cross gave prediction accuracy of ≈ 0.06, while 40 crosses with 10 lines per cross gave prediction accuracy of ≈ 0.075 (Figure). We assume that in both strategies different processes increase the accuracy with the addition of extra lines: In the first case, entire crosses were masked simulating the future prediction of an unphenotyped cross. In comparison, increasing the number of lines instead of number of crosses (while constraining the training set size) did not necessarily improve the prediction accuracy. The lines capture the crosses’ variance, and there will be a limit to how much more variance that additional lines will capture, hence no additional gain. The exception to this was adding fractions of the validation cross’ lines to the training set (Figure).

For item 2), we saw in Figure that using training sets comprised of exclusively unrelated crosses resulted in lower prediction accuracies than training sets that included related crosses. Using training sets comprised of either exclusively related crosses or related and unrelated crosses (half-and-half) both resulted in approximately the same prediction accuracy. The comparison between these three sets stops at about 800 lines in the training set, because beyond this point, additional crosses were no longer distinctively related or unrelated. Therefore, after this point the slope of increase in prediction accuracy is less steep, as the crosses added to the training set are less related.

For item 3), there was no observable connection between how well the cross could be predicted and the cross’ heritability or the observed phenotypic variance. Likewise, these values did not correspond to how well the data from the cross could be used to predict breeding values in other crosses.

One of the major practical implications of this study is that increased prediction accuracies can be obtained by balancing the training set for genomic selection with phenotypic and genomic data of multiple related crosses, which could be taken into account in advance when designing the training population, as previously proposed by Rincent et al., 2012. For existing data sets, a strategy may be applied of supplementing these with phenotypic data from previous trials (provided genotype-by-environment interaction is limited or can be accounted for by use of trait data for control lines). Although such data might be present within the context of a rolling breeding program, obtaining genomic data presents a bottleneck as this requires genotyping of (old) biological material that might not be readily available, and will require investment in at least low-density genotyping. In case high density genotype data sets are available for the parental lines, high density genotype information for their offspring populations can subsequently be obtained by imputation, as reported by Hickey et al. (2015), Gorjanc et al. (2017) and others.

### Conclusions

Genomic predictions of yield across 44 populations resulted in modest correlations between observed and predicted values. The correlations did increase with training set size, but by selecting training sets that comprised related crosses improved the correlation more than increasing training set size. The results also showed that if the training set size is fixed, using few lines from more crosses, rather than many lines from few crosses, resulted in higher correlations.

## Authors’ contributions

Wheat crosses were made by JL, EB, CB, PJ, SB, EF, BP, SS, CH; wheat yield trials were conducted by RJ, PH, EO and IJM; ARB co-ordinated genotyping; SME, RCG and GG performed data analysis; SME, JBB, RCG and JMH wrote the manuscript; JMH and IJM conceived the study, designed the experiment and led the project.

## Acknowledgements

The authors acknowledge the financial support from BBSRC project “GplusE: Genomic selection and Environment modelling for next generation wheat breeding” (grants BB/L022141/1 and BB/L020467/1) and the Medical Research Council (MRC) grant MR/M000370/1. The crosses, seed and genomic DNA for this study were contributed by KWS UK, RAGT Seeds Ltd., Elsoms Wheat Ltd and Limagrain UK.

## Competing interests

The authors declare that they have no competing interest.

## Supplementary materials

**Supplemental figure 1:**
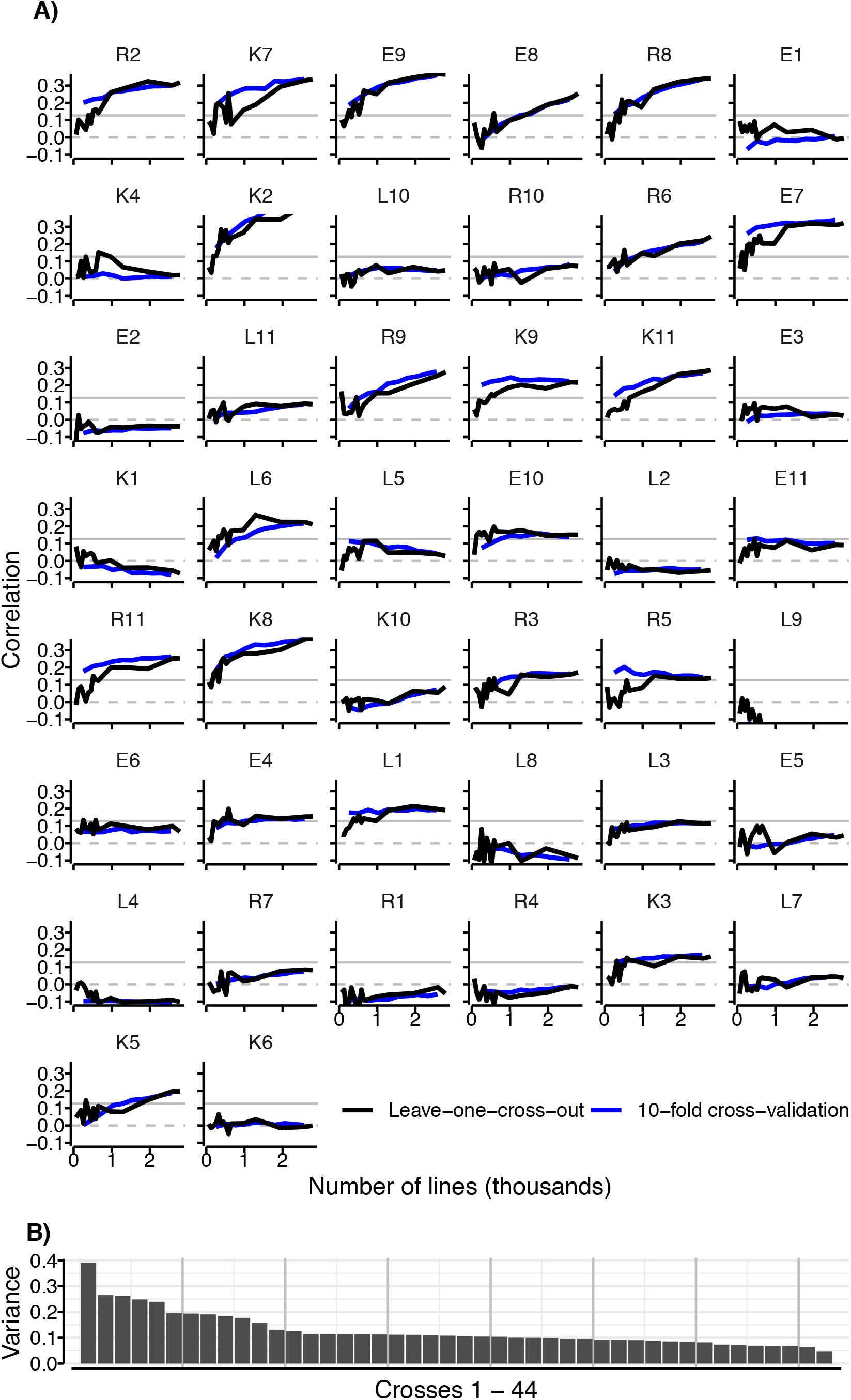
Per-cross correlation under two approaches. **(A), ordered by decreasing variance of crosses’ BLUEs (B).** Grey, horizontal lines are guides for zero correlation (dashed) and overall average correlation of 0.127 (solid). Crosses in A) are ordered with decreasing variance of their BLUEs, same order as in B).

